# 16S rRNA Modifications Are Dispensable for Viability but Collectively Optimize Ribosome Biogenesis and Translation Initiation in *Escherichia coli*

**DOI:** 10.64898/2026.09.08.748864

**Authors:** Natalie Åkesson, Anna af Klercker, Maheshwaran Sivakumar, Mikhail Metelev, Anna Knöppel, Magnus Johansson, Gerrit Brandis

**Affiliations:** Department of Cell and Molecular Biology, Uppsala University, Uppsala, Sweden; Uppsala Antibiotic Center, Uppsala University, Uppsala, Sweden

## Abstract

Ribosomal RNAs contain numerous conserved nucleotide modifications, yet the functional importance of most of these modifications remains unclear. In *Escherichia coli*, deletion of individual 16S rRNA modification enzymes generally produces only minor phenotypes, raising questions about their biological significance. Here, we generated a comprehensive collection of deletion mutants lacking individual and combined 16S rRNA modifications, culminating in a strain lacking all known 30S ribosomal subunit modifications. Despite the absence of all known 16S rRNA modifications, cells remained viable, exhibiting a fitness defect of ∼30% at 37 °C that increased to ∼50% at 20°C, consistent with impaired ribosome biogenesis. We identified strong epistatic interactions between modifications in the 3ʹ major and 3ʹ minor domains of 16S rRNA, resulting in disproportionately large effects on both fitness and antibiotic susceptibility. Live-cell single-molecule tracking revealed a marked increase in the fraction of non-translating ribosomes and a prolonged time required to enter productive translation, whereas translational elongation by actively engaged 70S ribosomes remained largely unaffected. In addition, fluorescence-based measurements showed that unmodified ribosomes exhibited increased stringency during translation initiation, reducing utilization of near-cognate start codons. Together, these findings demonstrate that 16S rRNA modifications are not essential for viability but collectively enhance the efficiency, robustness, and fidelity of ribosome assembly and translational initiation.

## Introduction

Ribosomes are essential molecular machines responsible for protein synthesis in all living cells. Their structural and functional features are highly conserved across Bacteria, Archaea, and Eukarya, underscoring their central role in cellular life.^1^ In *Escherichia coli*, the ribosome consists of three rRNAs (5S, 16S, and 23S) and 54 ribosomal proteins, organized into the 30S subunit comprised of the 16S rRNA and 21 proteins, and the 50S subunit comprised of 5S rRNA, 23S rRNA, and 33 proteins. During translation, these subunits associate to form the functional 70S ribosome.^2^ Beyond their primary sequences, ribosomal RNAs undergo extensive chemical modifications. In *E. coli*, 16S rRNA contains 11 modified nucleotides and 23S rRNA contains 25, which cluster near functional domains.^3,4^

Ribosome assembly is a highly structured yet dynamic process that can proceed through multiple parallel pathways.^5-7^ Specific nucleotide modifications are introduced at defined stages of assembly, suggesting a functional role in ribosome biogenesis.^8^ These modifications have been proposed to influence base stacking interactions that stabilize functionally active RNA conformations and/or act as check-points during subunit assembly.^9^ Indeed, modifications within the peptidyl transfer centre have been shown to be essential for *in vitro* assembly and functionality of 23S rRNA, and deletion of specific modification enzymes leads to accumulation of assembly intermediates and reduced translational fidelity *in vivo*.^10-16^ However, subsequent studies demonstrated that unmodified 23S rRNA can fold into catalytically active 50S particles *in vitro* with the help of osmolytes.^17^ Similarly, *in vitro* synthesized 16S rRNA can be assembled into functional 30S subunits.^18^ Furthermore, no single modification enzyme is essential for growth in *E. coli*, and several studies have shown that the deletion of individual modification enzymes has minimal effects *in vivo*, with the notable exception of *rlmE*.^10,19,20^ These findings challenge the presumed necessity and functional importance of nucleotide modifications in the *E. coli* ribosome.

A possible explanation for this discrepancy is the existence of multiple parallel pathways for ribosome assembly.^5-7^ Deletion of an individual modification enzyme may disrupt specific pathways, yet overall assembly remains largely unaffected due to redundancy in the assembly process. To investigate this possibility, *E. coli* strains carrying multiple modification enzyme deletions have been constructed.^12,15,21-24^ Remarkably, removal of all seven pseudouridine synthetases had only minimal effects *in vivo*.^24^ For the 16S rRNA, three double deletions (Δ*rsmAJ*, Δ*rsmBD*, and Δ*rsmHI*) have been examined, with Δ*rsmHI* being the only combination that significantly impaired cell viability and translational fidelity.^12,21,22^ In the case of 23S rRNA, deletion combinations have primarily targeted modifications within the peptidyl transferase center.^15,23,25^ These studies revealed synergistic interactions between specific mutations, most notably that the absence of modifications by RlmB, RlmL, RlmN, or RluC amplify the detrimental impact of losing RlmE. Collectively, these findings underscore the complexity of modification networks and highlight that systematic analysis of cumulative effects from multiple deletions is essential to fully elucidate the functional interplay of rRNA modifications in ribosome assembly and performance.

A complicating factor for such systematic studies is that recent attempts to combine modification enzyme deletions have relied on transductions from strains of the KEIO collection.^15,23-25^ Although this approach is fast and efficient for introducing individual gene deletions, it poses significant risks when generating multiple knockouts. The KEIO deletions are constructed using a kanamycin resistance cassette flanked by FRT sites.^26^ This design enables sequential introduction of deletions via transduction, followed by FLP-mediated recombination to excise the resistance marker, which allows subsequent rounds of deletion. However, each excision leaves behind a FRT scar. These scars serve as recognition sites for FLP recombinase and can lead to unwanted chromosomal rearrangements. Directly oriented scars may cause duplication or deletion of the intervening sequences, whereas inverted orientations can result in inversions.^27,28^ Consequently, as FRT scars accumulate, genome stability can become severely compromised. In addition, KEIO deletions were designed for high-throughput construction and remove entire coding sequences except for the last 21 nucleotides, which may include ribosome-binding sites for downstream genes.^26^ Importantly, several modification enzyme genes harbor promoters and regulatory elements for adjacent genes within their coding sequences.^29^ Deletion of these regions can introduce confounding polar effects, complicating interpretation of phenotypes. Together, these factors underscore the need for careful design and interpretation of multiple-deletion experiments, as observed phenotypes may reflect unintended genomic rearrangements or regulatory disruptions rather than the absence of rRNA modifications alone.

In this study, we investigated the functional consequences of removing modifications within the 30S ribosomal subunit of *E. coli*. Using careful design and scar-free recombineering methods, we created a set of strains carrying individual modification enzyme deletions and combined these mutations according to the functional domains of 16S rRNA to generate strains with progressively under-modified and ultimately modification-free 30S subunits. We find that strains carrying fully unmodified 30S subunits remain viable, exhibiting only a ∼30% reduction in fitness at 37 °C. This fitness defect became more pronounced at lower temperatures, consistent with impaired ribosome assembly. Furthermore, we identified epistatic interactions among modification enzymes acting in the 3’ major and 3’ minor domains of 16S rRNA, affecting both fitness and antibiotic susceptibility. Finally, we show that unmodified 30S subunits display increased stringency during translation initiation, leading to reduced utilization of non-AUG start codons, while also increasing the fraction of non-translating ribosomes approximately fivefold. Together, these findings demonstrate that 16S rRNA modifications are not required for viability but collectively promote ribosome assembly, translational capacity, and initiation fidelity.

## Results

### Construction and Combination of rRNA Modification Enzyme Deletions

Scar-free deletions of modification enzyme genes were designed following the same basic principles as the KEIO collection.^26^ To avoid disrupting ribosome binding sites for downstream genes, the last 21 nucleotides of each coding sequence were retained. Likewise, the first 18 nucleotides, including the start codon, were preserved. This design ensures that the deletion results in the translation of a short peptide of 12 amino acids instead of the original enzyme, which minimizes polar effects on operons (Fig. 1A). Three of the genes (*rsmA*, *rsmH*, and *rsmI*) contain known regulatory sites for neighbouring genes.^29^ For these modification enzymes, the deletions were designed to preserve all identified regulatory elements (Fig. 1B). In two of these cases (*rsmA* and *rsmH*) a significant proportion (>40%) of the genes remain intact, but the deletions remove the SAM-binding motifs essential for methylation activity (Supplementary Table S1).^30,31^

**Fig. 1.**
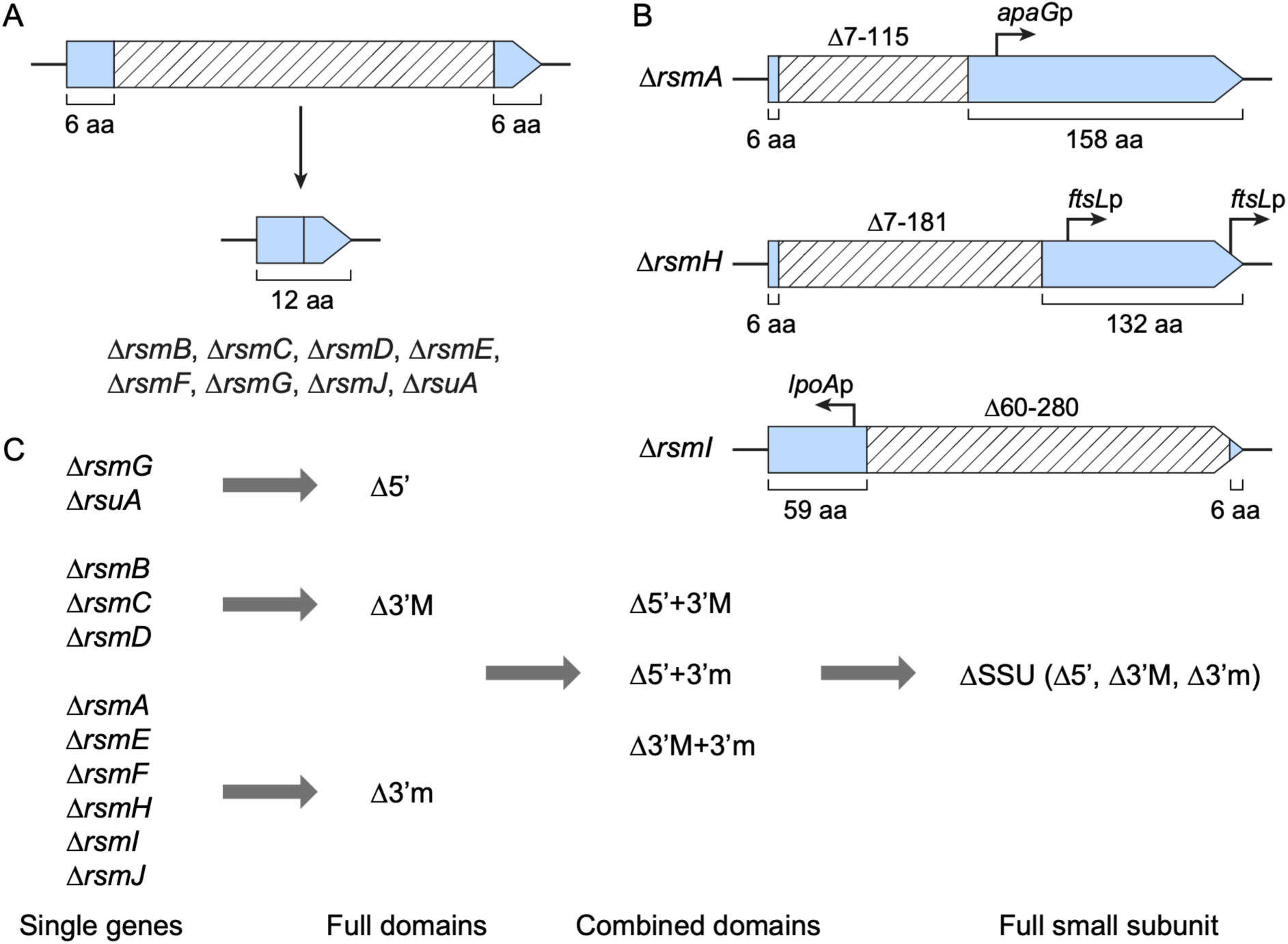
Overview of strain constructions. (**A**) Deletions were generally designed to maintain the first and last six amino acids resulting in the translation of a 12 amino acids long peptide. (**B**) The *rsmA*, *rsmH*, and *rsmI* genes were only partially deleted to preserve regulatory elements for neighbouring genes. (**C**) The deletions of modification enzymes were combined to generate strains with unmodified domains, combinations of unmodified domains, and fully unmodified small ribosomal subunits.

Deletions of modification enzyme genes were grouped according to the domain structure of the small ribosomal subunit.^32^ The deletions of the two enzymes that modify the 5’ domain (Δ*rsmG* and Δ*rsuA*) were combined to generate a strain with an unmodified 5’ domain (Δ5’). Similarly, Δ*rsmB*, Δ*rsmC*, and Δ*rsmD* were combined to create a strain lacking modifications in the 3’ major domain (Δ3’M); and Δ*rsmA*, Δ*rsmE*, Δ*rsmF*, Δ*rsmH*, Δ*rsmI*, and Δ*rsmJ* were combined to produce a strain with an unmodified 3’ minor domain (Δ3’m).

These single-domain modification deletion strains were further combined to generate strains with double-domain modification deletions (Δ5’+3’M, Δ5’+3’m, and Δ3’M+3’m) and a strain with a fully unmodified small subunit (ΔSSU) (Fig. 1C). All strains carrying multiple gene deletions were subjected to whole-genome sequencing to verify the intended deletions. No significant secondary mutations arose during the strain construction (Supplementary Table S2). In total, eighteen strains containing between one and eleven modification enzyme deletions were constructed for subsequent analyses.

### Non-Additive Fitness Costs Indicate Defects in Under-Modified Ribosomes

Previous studies have reported modest growth defects for strains lacking individual small- subunit rRNA modification enzymes, although the results have often been inconsistent across laboratories.^10,12,21,33^ Moreover, several modification gene deletions were found to exhibit more pronounced fitness costs in competitive growth assays than during measurement of exponential growth rates alone.^34,35^ To resolve these discrepancies, we quantified both exponential growth rates and competitive fitness across our panel of eighteen strains at 20 °C, 30 °C, 37 °C, and 42 °C (Fig. 2, Supplementary Fig. S1, and Supplementary Tables S3-S5).

**Fig. 2.**
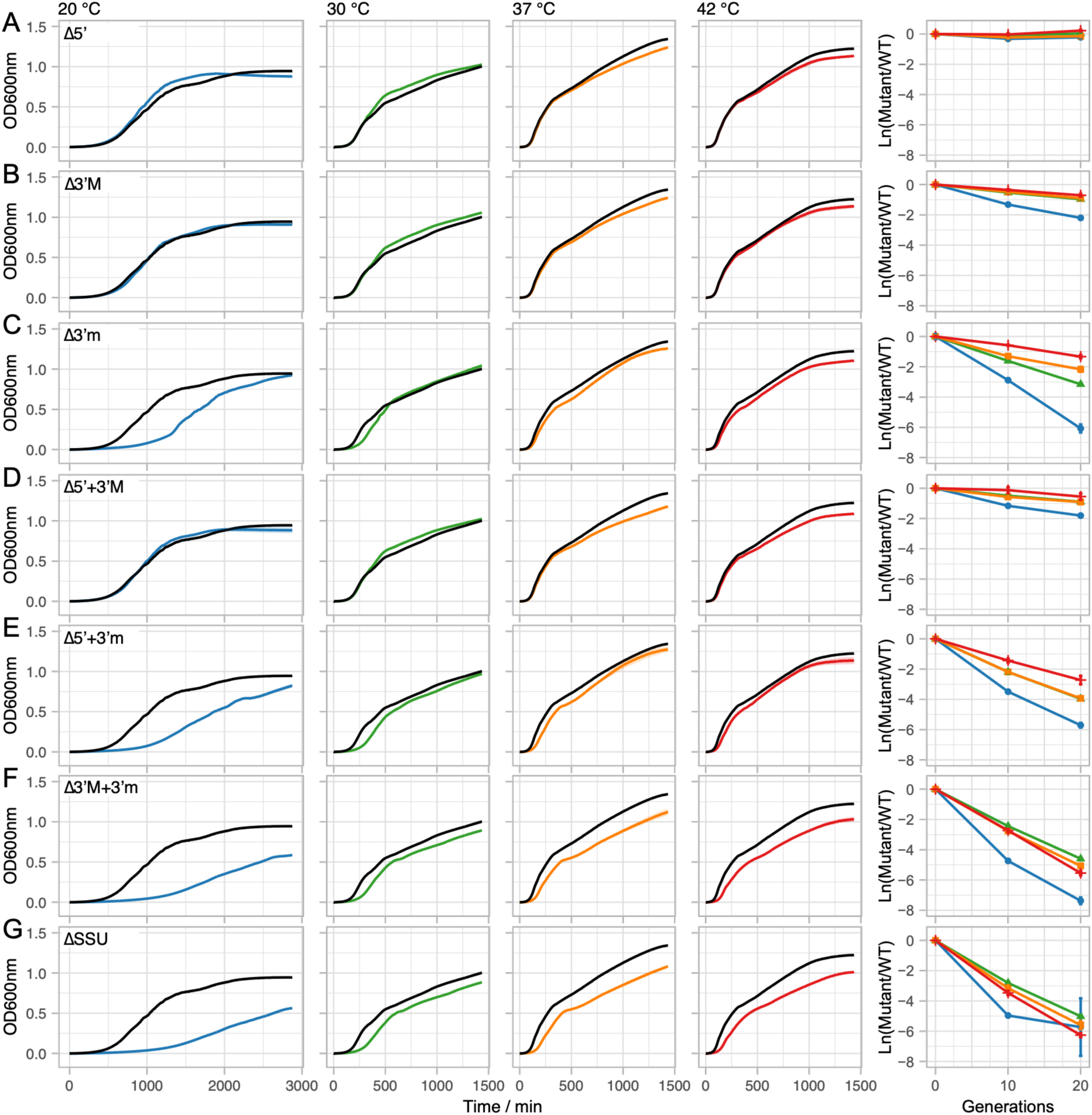
Analysis of growth characteristics. (**A** - **G**) Growth curves and competition experiments of isolates with unmodified domains and combinations thereof. Strain genotypes are indicated in each panel and growth temperatures are shown at the top of the growth curves. Growth curves of the constructed strains are shown in blue (20 °C), green (30 °C), orange (37 °C), and red (42 °C) and the growth curves of the isogenic wild type at each temperature are shown in black. All growth curves are the average of three biological replicates with standard variation shown as shaded region along curve. Competitions were performed at 20 °C (blue, circles), 30 °C (green, triangles), 37 °C (orange, squares), and 42 °C (red, line). All competition data represent the average ± standard deviation of three biological replicates. Values were normalized to account for the fitness cost of the fluorescence marker and differences in initial starting ratios of the competitors.

The competition experiments reveal that six single-gene deletion strains (Δ*rsmC*, Δ*rsmE*, Δ*rsmF*, Δ*rsmG*, Δ*rsmJ*, and Δ*rsuA*) show no detectable fitness cost at any of the measured temperatures. The remaining five deletions (Δ*rsmA*, Δ*rsmB*, Δ*rsmD*, Δ*rsmH*, and Δ*rsmI*) impose a significant fitness cost of 4 – 14% per generation at 20 °C, which progressively diminishes as temperature increases. At 30 °C, only three deletions (Δ*rsmA*, Δ*rsmD*, and Δ*rsmH*) retain measurable costs (3 – 6% per generation), and at 37 °C only Δ*rsmA* and Δ*rsmD* remain costly (4% per generation). At 42 °C, a significant (3% per generation) defect persists only for Δ*rsmA* (Supplementary Fig. S1, Supplementary Table S5). A similar temperature dependence is observed in strains lacking modifications of entire domains or combinations thereof. The Δ5’ strain is the only multi-deletion strain without a detectable fitness defect at any temperature. All other compound-deletion strains show substantial fitness losses, decreasing from 9 – 50% per generation at 20 °C to 3 – 31% at 42 °C (Fig. 2, Supplementary Table S5). These temperature-dependent costs are consistent with ribosome assembly defects in under-modified small subunits^15,36,37^ and increase with the number of missing modifications. Fitness estimates based on exponential growth rates follow the same general pattern but often indicate larger defects than competition assays (Fig. 2, Supplementary Fig. S1, and Supplementary Tables S3-S4), suggesting that the assembly-associated limitations are more pronounced during rapid growth.

Notably, at every temperature tested, the competitive fitness cost of the double non-modified domain strain Δ3’M+3’m is significantly greater (17.3% ± 7.6%, P = 0.023, paired two-sided t-test) than predicted by an additive model based on the individual Δ3’M and Δ3’m strains. Furthermore, introducing deletions affecting the 5’ domain to generate the ΔSSU strain imposes an additional 2 – 5% cost per generation, even though the Δ5’ strain alone shows no measurable defect (Supplementary Table S5). Together, these results indicate that loss of rRNA modifications leads to defects in ribosome assembly and that these defects exhibit strong epistatic interactions when multiple domains are unmodified.

### Epistatic Interactions Between 16S rRNA Modifications Alter Antibiotic Susceptibility

The lack of methylation at position A1519 of the 16S rRNA by RsmA results in increased resistance to the antibiotic kasugamycin.^38^ Deletions of *rsmA*, *rsmF*, or *rsmG* have also been linked to decreased susceptibility to various other aminoglycoside antibiotics. However, the changes in susceptibility were generally minimal *(e.g.* 2-fold change in MIC) and within the natural variation of the assays.^39-42^ We tested susceptibility levels of our strains against a set of twelve ribosome-binding antibiotics (kasugamycin/KSG, gentamicin/GEN, streptomycin/STR, amikacin/AMK, spectinomycin/SPT, tetracycline/TET, tigecycline/TGC, linezolid/LZD, chloramphenicol/CHL, erythromycin/ERY, clindamycin/CLI, and tiamulin/TIA) and one non-ribosomal targeting antibiotic (rifampicin/RIF) as control to identify if the lack of modification genes result in a significant change in susceptibility (Fig. 3). We decided to start with the ΔSSU strain that carries a fully unmodified small ribosomal subunit since the effect on antibiotic susceptibility is expected to be highest in this strain. We found significantly elevated susceptibility to five of the tested antibiotics (STR, AMK, TET, ERY, and TIA) and increased resistance to one antibiotic (KSG). To identify which of the modification enzyme deletions contribute to these changes, we measured antibiotic susceptibilities in strains that carry unmodified domains (Δ5’, Δ3’M, and Δ3’m) and combinations thereof (Δ5’+3’M, Δ5’+3’m, and Δ3’M+3’m). In agreement with the previous data, none of the strains displayed changed susceptibility to the seven antibiotics (GEN, SPT, TGC, LZD, CHL, CLI, and RIF) that were also unchanged in the ΔSSU strain (Fig. 3). Increased resistance to kasugamycin was caused by modification enzyme deletion in the 3’m domain that include *rsmA*.^38^ For the other antibiotics (STR, AMK, TET, ERY, and TIA), a combination of modification enzymes deletions from the 3’M and 3’m domain (Δ3’M+3’m) was necessary and sufficient for the observed increased susceptibilities in the ΔSSU strain (Fig. 3). These data indicate epistatic interactions between the modifications of domain 3’M and 3’m which agrees with the epistatic effects between these two domains in the competitive fitness measurements.

**Fig. 3.**
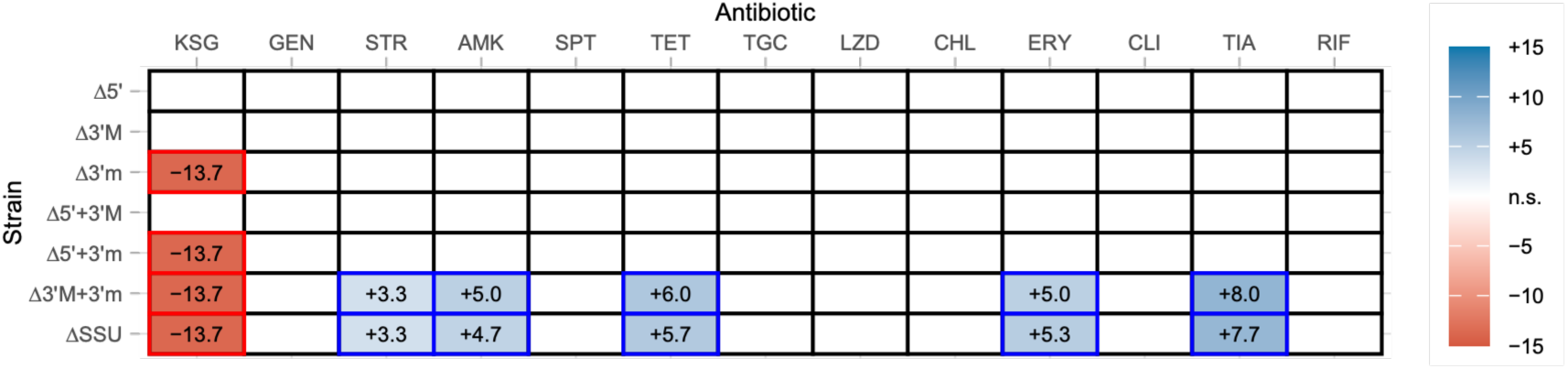
Antibiotic susceptibility testing. Change in the zone of inhibition between wild-type *E. coli* and strains carrying under-modified ribosomes. Increased antibiotic susceptibility is indicated in blue, and increased resistance in red. Empty fields represent no significant change. All values are averages of three biological replicates. See Supplementary Data 1 for all individual measurements.

### Loss of 16S rRNA Modifications Increases the Stringency of Translation Initiation

Functional ribosomes are essential for translational fidelity.^43,44^ To determine whether loss of 16S rRNA modifications affects translational initiation fidelity, we constructed a dual-fluorescence reporter system that measures initiation from non-AUG start codons (GUG, UUG, and CUG) (Fig. 4A). The reporter includes a constitutively expressed yellow fluorescent protein (*yfp*) that serves as an internal control, allowing normalization of blue fluorescent protein (*bfp*) expression and compensating for variations in mRNA stability that may arise when reduced translation leads to increased transcript degradation. Comparison of *bfp* expression from AUG and non-AUG start codons in the wild-type strain showed that GUG and UUG support expression at 41% and 21% of AUG levels, respectively (Fig. 4B). In the ΔSSU strain, initiation from both non-AUG codons was significantly reduced. Expression from the GUG start codon decreased to 32% (P = 1.243 × 10^-6^, two-sided t-test), while expression from the UUG start codon decreased to 12% (P = 1.696 × 10^-4^, two-sided t-test). No detectable *bfp* expression was observed from the CUG start codon in either strain. Together, these results indicate that loss of 16S rRNA modifications increases the stringency of translation initiation, resulting in reduced utilization of near-cognate start codons.

**Fig. 4.**
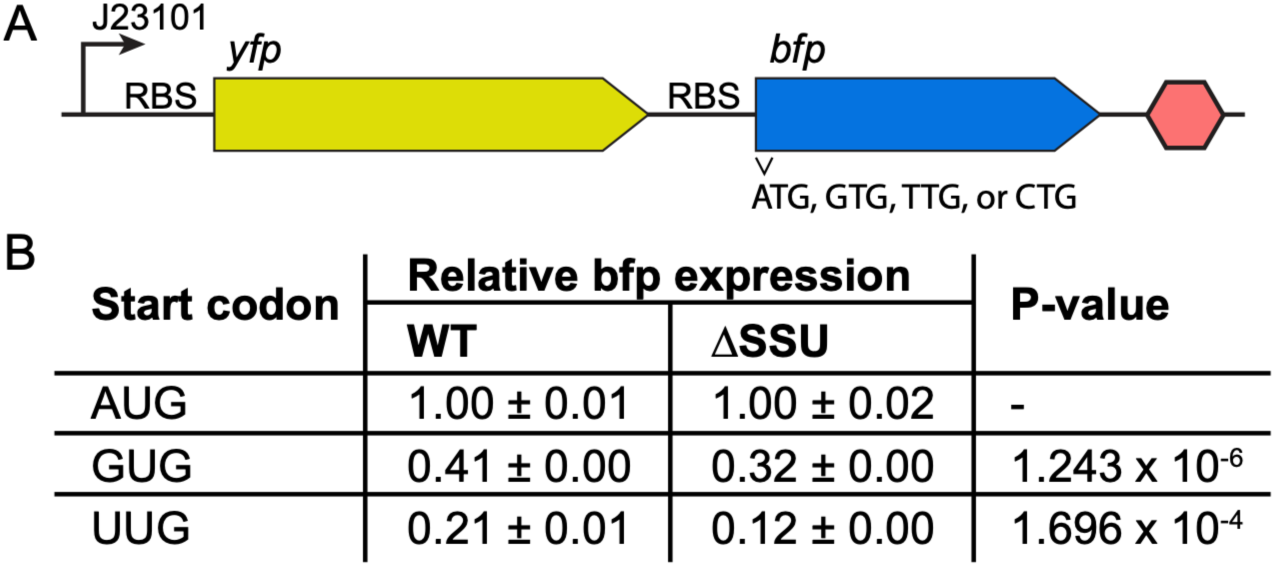
Translation initiation fidelity assay. (**A**) Schematic of the dual-fluorescence reporter system. Transcription of *yfp* and *bfp* is driven by the constitutive promoter J23101. Each gene contains an independent ribosome-binding site. Translation of *yfp* is initiated exclusively from an AUG start codon and serves as an internal control, whereas *bfp* carries either an AUG, GUG, UUG, or CUG start codon to measure initiation from cognate and near-cognate start codons. (**B**) Relative *bfp* expression from different start codons in wild-type *E. coli* and the ΔSSU strain, normalized to *yfp* expression. Values represent mean ± standard deviation of three biological replicates. No detectable *bfp* expression was observed from the CUG start codon in either strain.

### Loss of 16S rRNA Modifications Increases the Fraction of Non-Translating Ribosomes

Next, we performed live-cell single-molecule tracking of fluorescently labelled ribosomes to determine how the absence of 16S rRNA modifications affects the *in vivo* kinetics of translation initiation and elongation. For this purpose, wild-type *E. coli* and the ΔSSU strain were transformed with plasmid-encoded *rrnB* operons carrying MS2 aptamer insertions in either the 16S or 23S rRNA, enabling specific fluorescent labelling using a co-expressed MS2 coat protein-HaloTag fusion. Because the chromosomal rRNA operons remained intact, only the subset of ribosomes assembled from the plasmid-derived rRNAs carried the MS2 aptamer and could be specifically labelled and tracked.

Tracked ribosomal subunits were classified into two diffusion states corresponding to free ribosomes (fast diffusion state) and ribosomes engaged in translation (slow diffusion state), as previously described.^45^ In the ΔSSU strain, the steady-state occupancy of the fast diffusion state increased substantially for both subunits. For 30S subunits, the fraction of ribosomes in the fast diffusion state increased 4.5-fold from 5.6% to 25.6%, while for 50S subunits it increased 3.8-fold from 6.9% to 26.4% (Fig. 5A). Consistent with this observation, the average dwell time in the fast diffusion state increased from 1.1 s to 5.2 s for 30S subunits and from 1.1 s to 4.9 s for 50S subunits (Fig. 5B). These results indicate that loss of 16S rRNA modifications substantially prolongs the time required for ribosomes to enter productive translation cycles. In contrast, only minor changes were observed in the dwell time of the slow diffusion state. For 30S subunits, the average dwell time decreased from 22.7 s to 16.9 s, whereas for 50S subunits it changed from 20.1 s to 21.4 s (Fig. 5C). These differences were within the variation of the assay and did not reveal a consistent effect on elongation.

**Fig. 5.**
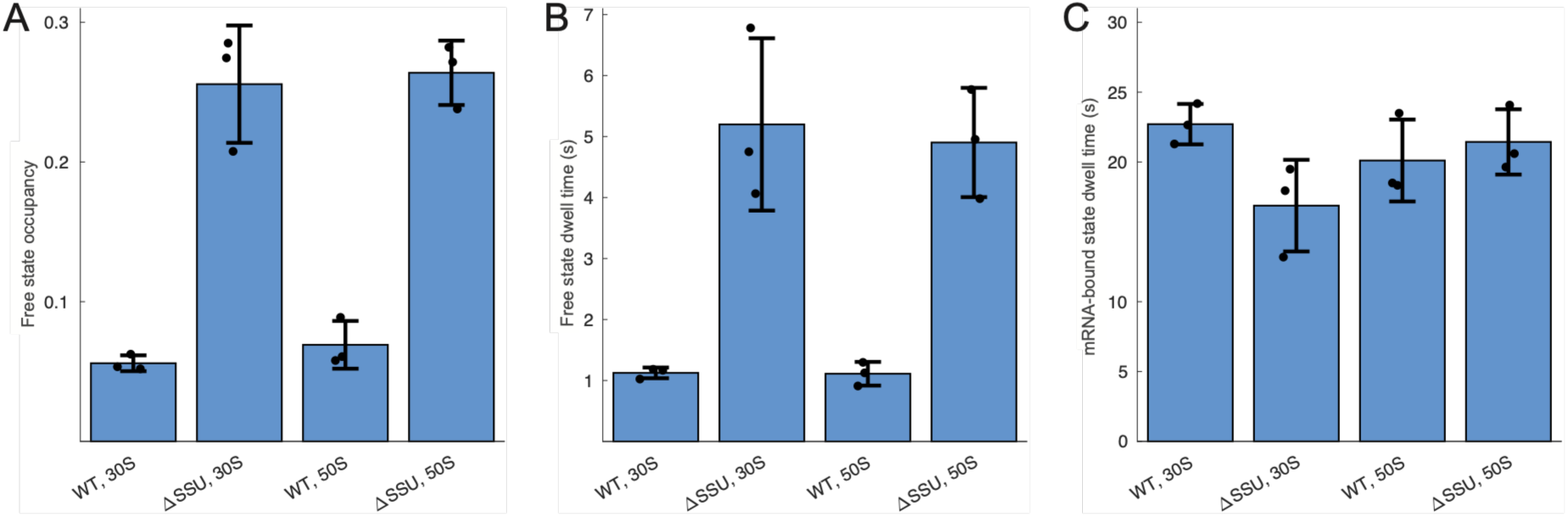
Estimated occupancy and dwell times of ribosomal subunits. (**A**) Occupancy of the fast diffusion state, representing freely diffusing ribosomal subunits, for 30S (16S rRNA) and 50S (23S rRNA) subunits in wild-type *E. coli* and the ΔSSU strain. Dwell times of 30S and 50S ribosomal subunits in wild-type *E. coli* and the ΔSSU strain in (**B**) the fast diffusion state (free ribosomes) and (**C**) the slow diffusion state (mRNA-bound ribosomes). Bar plots show the results from coarse-grained 6-state HMM models. Values represent mean ± standard deviation of three biological replicates. Individual points represent independent microscopy experiments.

Together, these data indicate that loss of 16S rRNA modifications primarily impairs mRNA binding rather than translation by actively engaged ribosomes. As a consequence, the fraction of ribosomes engaged in translation decreases from approximately 94% in the wild type to 74% in the ΔSSU strain. These results indicate that 16S rRNA modifications primarily promote formation of initiation-competent ribosomal subunits, while having little or no effect on translational elongation once 70S ribosomes have entered productive translation.

## Discussion

The functional importance of 16S rRNA modifications has remained difficult to define despite decades of investigation. Comparison between previous studies has been complicated by the use of different *E. coli* K-12 backgrounds, most commonly BW25113 and MG1655, which differ at numerous loci and can display strain-specific phenotypes.^10,26,34,46^ Furthermore, analyses of multi-deletion strains have frequently relied on derivatives of the KEIO collection, introducing the possibility of confounding effects from FRT scars, chromosomal rearrangements, and disruption of regulatory elements controlling neighbouring genes. ^26-29^ In addition, competitive fitness measurements have previously relied on selection markers derived from the kanamycin resistance cassette itself.^21^ Because this cassette imposes a measurable fitness cost, interpretation of such experiments can be challenging.^34^ The scar-free strain collection generated here overcomes these limitations and provides a systematic framework for analysing the cumulative effects of 16S rRNA modification loss in a single genetic background.

Among the individual modification enzymes examined, RsmA emerged as the most important contributor to cellular fitness. Deletion of *rsmA* resulted in measurable fitness costs across all temperatures tested and contributed prominently to the phenotypes observed in strains carrying multiple modification defects (Supplementary Table S5). This finding is consistent with extensive evidence that RsmA functions as a late-stage ribosome biogenesis factor rather than simply a methyltransferase. Structural studies in *Thermus thermophilus* demonstrated that dimethylation of A1518 and A1519 stabilizes interactions between helices 44 and 45 of the decoding centre.^47^ Cryo-EM analyses in *E. coli* further suggest that methylation by RsmA promotes conformational rearrangements required for final maturation of the 30S subunit and efficient processing of 17S precursor rRNA to mature 16S rRNA.^48^ Consistent with this model, deletion of *rsmA* impairs 17S processing at low temperature, interferes with displacement of the assembly factor RbfA by IF3, and increases initiation from non-canonical start codons.^10,11,30,49^ More recently, RsmA has been proposed to function in a quality-control pathway that rescues inactive mature-like 30S particles through targeted remodelling and remethylation.^50^ Together, these findings provide a mechanistic explanation for why loss of RsmA exerts a substantially larger effect than most other 16S rRNA modification enzymes. The strong temperature dependence of the Δ*rsmA* phenotype (Supplementary Fig. S1, and Supplementary Tables S3-S5) further supports a role during ribosome biogenesis, as defects in rRNA folding and maturation are generally exacerbated at lower temperatures.^15,36,37^

The strong epistatic interactions observed between modifications in the 3ʹ major and 3ʹ minor domains suggest that the functional importance of individual modifications is buffered by the broader modification network. Simultaneous loss of modifications from both domains produced substantially larger fitness defects and antibiotic susceptibility changes than expected from the effects of the individual domain deletions alone. Similar synergistic interactions have previously been reported for combinations of 23S rRNA modification defects.^15,23,25^ These findings support a model in which rRNA modifications contribute collectively to ribosome assembly, such that disruption of a single modification can often be accommodated through alternative assembly routes, whereas removal of multiple modifications progressively reduces assembly robustness and the production of functional ribosomes.

RsmF represents a special case among the enzymes examined because it modifies both 16S rRNA and tRNA substrates (Supplementary Table S1). Despite this dual specificity, deletion of *rsmF* produced no detectable fitness defect under any growth condition tested (Supplementary Fig. S1, and Supplementary Tables S3-S5). This suggests that the absence of RsmF-dependent tRNA modifications contributes little, if at all, to the phenotypes reported here. Consequently, the defects associated with the ΔSSU strain are most likely explained by loss of 16S rRNA modifications rather than secondary effects arising from altered tRNA modification.

The dual-fluorescence reporter experiments revealed that ribosomes lacking 16S rRNA modifications exhibit increased stringency during translation initiation, resulting in reduced utilization of near-cognate start codons (Fig. 4). This phenotype is likely to have direct physiological consequences because approximately 17.5% of annotated *E. coli* genes initiate with GUG or UUG start codons rather than the canonical AUG codon (14.3% GUG and 3.2% UUG).^46^ Reduced expression of this substantial subset of genes is likely to contribute directly to the fitness defects observed in under-modified ribosomes.

The single-molecule tracking experiments demonstrate that the primary consequence of removing 16S rRNA modifications is a reduction in the efficiency with which ribosomes enter productive translation cycles (Fig. 5A and B). In contrast, the average dwell time of translating ribosomes was largely unaffected (Fig. 5C). These observations suggest that the major function of 16S rRNA modifications is to promote formation of initiation- competent ribosomal subunits rather than to optimize the catalytic performance of actively translating 70S ribosomes. This interpretation agrees with structural and genetic studies that place many 16S rRNA modifications within assembly pathways rather than directly within the elongation cycle.^8-11^

Several limitations should nevertheless be considered. First, the current single-molecule tracking framework reports average dwell times for the ribosome population and therefore cannot distinguish between a model in which all ribosomes initiate more slowly and one in which a subset of ribosomes fails to initiate translation entirely. Second, particles that become severely misassembled and are rapidly degraded are unlikely to be detected by the tracking approach. Consequently, the measured decrease in the fraction of translating ribosomes may underestimate the full impact of modification loss on ribosome biogenesis. Future studies that directly quantify assembly intermediates and catalytic activity of unmodified ribosomes should help to resolve these possibilities.

Collectively, our results demonstrate that 16S rRNA modifications are not required for viability but nevertheless provide substantial fitness advantages by optimizing ribosome biogenesis and translational capacity. The viability of cells lacking all known 30S subunits modifications reveals an unexpected robustness of the bacterial translation machinery. At the same time, the strong epistatic interactions observed between modifications in different 16S rRNA domains indicate that these modifications function as an interconnected network rather than as independent structural elements. The pronounced fitness defects, altered antibiotic susceptibilities, increased initiation fidelity, and accumulation of non-translating ribosomes observed in the ΔSSU strain suggest that individual assembly defects can often be buffered, whereas the combined loss of multiple modifications progressively reduces the efficiency and robustness of ribosome biogenesis. Rather than acting as essential functional elements, 16S rRNA modifications appear to serve as evolutionary refinements that maximize the performance, reliability, and assembly efficiency of the ribosome.

## Methods

### Bacterial strains and growth conditions

Strains within this study were derived from *Escherichia coli* K-12 strain MG1655 and are listed in Supplementary Table S2.^46^ Bacteria were grown in Luria Broth (LB; 10 g L^-1^ tryptone, 5 g L^-1^ yeast extract, 5 g L^-1^ NaCl) or on LB agar plates (LA; LB solidified with 1.5% Oxoid agar). Tetracycline (15 mg L^-1^), chloramphenicol (25 mg L^-1^), and sucrose (50 g L^-1^) were added to the media as required.

### Generation and combination of gene deletions

Genetic engineering was performed using λ-Red recombineering with the temperature- sensitive pSIM5-tet plasmid. ^51^ In general, deletions were designed to remove all codons except the first six and last six, resulting in a short 12-amino-acid peptide to minimize polar effects on downstream genes. For genes with known regulatory elements, deletions were adjusted accordingly (Fig. 1, Supplementary Table S1). Scar-free gene deletions were constructed using DiRex^52^, except for *rsmI*, where DiRex primers failed to amplify the required resistance cassette. In this case, the counter-selectable Acatsac1 cassette was used in combination with ssDNA recombineering to generate a scar-free deletion.^51,53,54^ Deletions were combined by P1 transduction using DiRex intermediates.^52^ Dup-In was employed to transfer the scar-free *rsmI* deletion.^55^ All oligonucleotides used for deletion construction are listed in Supplementary Table S6.

### Local and whole genome sequencing

All PCRs for sequence confirmation of deletions and mutations were performed using 2x Taq Master Mix (Thermo Scientific) according to the manufacturer’s instructions. PCRs followed the standard protocol: 95 °C for 5 min; 35 cycles of 95 °C for 30 s, 50 °C for 30 s, and 72 °C for 3 min; followed by a final extension at 72 °C for 10 min. Fragment sizes were verified on a 1% agarose gel run for 30 min at 100 V, and purified PCR products were sequenced at Macrogen (Denmark). All primers used for amplification and sequencing are listed in Supplementary Table S6. For whole-genome sequencing, chromosomal DNA was extracted using the UltraClean Microbial Kit (Qiagen) and sequenced at Macrogen (Denmark). Sequence analyses were performed using CLC Genomics Workbench version 25.0.3. Raw whole-genome sequencing data are available in the NCBI SRA database.

### Exponential growth rate measurements

Pre-cultures of each strain were grown overnight with shaking at 37 °C. Cultures were diluted 1,000-fold in fresh LB, and 300 µL was loaded into each well of a Bioscreen Honeycomb 2 plate. Cultures were incubated under continuous shaking for 48 h at 20 °C, or for 24 h at 30 °C, 37 °C, or 42 °C in a Bioscreen C Pro instrument, and OD_600nm_ was recorded every 5 min. Exponential growth rates were calculated by fitting an exponential curve over the initial 0.1 increase in OD_600nm_. Measurements were performed in three biological replicates and significance was determined using a two-sided t-test comparing the doubling times of the mutant strains with those of the isogenic wild type. All raw growth curve data are provided in Supplementary Data 2.

### Competition experiments

Competitors were labeled with chromosomal *yfp* or *bfp* fluorescence genes inserted at the *galK* locus.^56^ Strains carrying deletions of modification enzyme genes were compared to their isogenic wild-type counterparts. Overnight cultures of both competitors were grown independently in LB broth with shaking at 37 °C. Cultures were then mixed at a 1:1 ratio (T_0_ sample), diluted 1,000-fold into fresh LB, and incubated overnight with shaking at 20 °C, 30 °C, 37 °C, or 42 °C allowing approximately 10 generations of growth (T_1_ sample). This dilution and overnight growth cycle was repeated once more (T_2_ samples). The ratio of *yfp*- to *bfp*- labeled cells in each sample was determined by counting 30,000 cells using a MACS cell separation system (Miltenyi Biotec). Selection coefficients were calculated using a regression model as previously described.^57^ T_2_ samples where one of the competitors accounted for less than 1% of the counted cells were not used for the calculations. All experiments were performed in three biological replicates and a two-sided t-test was performed to test for significance. Fitness costs larger than 2% per generation with a P-value below 0.05 were deemed significant.

### Antibiotic susceptibility measurements

Antibiotic susceptibility was assessed using standard disc diffusion tests. Colonies were resuspended in 0.9% NaCl to a turbidity equivalent to 0.5 McFarland, and bacteria were spread onto LA plates using sterile cotton swabs. Antibiotic discs were applied, and plates were incubated for 18–20 h at 37 °C. All tests were performed in biological triplicates. Changes in the zone of inhibition were considered significant if the mean difference exceeded 2 mm and a two-sided t-test indicated statistical significance (P < 0.05). A complete list of tested antibiotics and the amount of antibiotic per disc is provided in Supplementary Table S7.

### Translation initiation fidelity assay

The dual-fluorescence reporter system was integrated into the *galK* locus of wild-type *E. coli* and the ΔSSU strain. Overnight cultures were grown in LB broth with shaking at 37 °C. Stationary-phase cultures were diluted 2,000-fold in PBS, and *yfp* and *bfp* fluorescence were measured for 30,000 cells using a MACSQuant Analyzer (Miltenyi Biotec). For each cell, mean *bfp* fluorescence was normalized to the corresponding mean *yfp* fluorescence to control for differences in reporter expression and mRNA abundance. The resulting normalized *bfp* values were averaged for each strain and expressed relative to those obtained with the reporter carrying an AUG start codon, which was set to 100%.

### Single-molecule tracking

The *rrnB* operons containing the MS2 aptamer inserted in h6 of 16S rRNA or H98 of 23S rRNA were PCR amplified from pAM552-rrnB-h6 and pAM552-*rrnB*-H98,^45^ respectively, using primers 5ʹ-AAATTGAAGAGTTTGATCATGGCTCAG-3ʹ and 5ʹ-AAGGTTAAGCCTCACGGTTCATT-3ʹ. The low-copy plasmid pSC101-P59-oASD-h6-MS2, previously used for tracking of ribosomal subunits,^45^ was PCR amplified using primers 5ʹ-TTGAGCTAACCGGTACTAATGAACCG-3ʹ and 5ʹ- GCCAGCGTTCAATCTGAGCCAT-3ʹ. The corresponding *rrnB* operon and plasmid fragments were assembled by Gibson Assembly, resulting in plasmids pSC101-P59-*rrnB*-h6 and pSC101- P59-*rrnB*-H98. These plasmids were transformed into wild-type *E. coli* and the ΔSSU strains.

Sample preparation, single-molecule tracking and microscopy data processing were performed as described previously for tracking of ribosomal subunits labeled via MS2 aptamer,^45^ with the exception of the optical setup. Imaging was performed on an inverted Nikon Ti2-E microscope equipped with a CFI Plan Apo Lambda 1.45/100× objective (Nikon), and phase-contrast and fluorescence images were acquired using an ORCA-Quest camera (Hamamatsu). Mini-colonies were imaged at 37 °C using stroboscopic laser illumination with a 3 ms laser pulse per 30 ms camera exposure using a 546 nm laser (2RU-VFL-P-2000-546- B1R 2000 mW, MPB Communication) with the power density of 3 kW/cm^2^ on the sample plane. Single-molecule trajectories were constructed and analyzed using the previously described pipeline and HMM approach.^45^

---

**Fig. S1.**
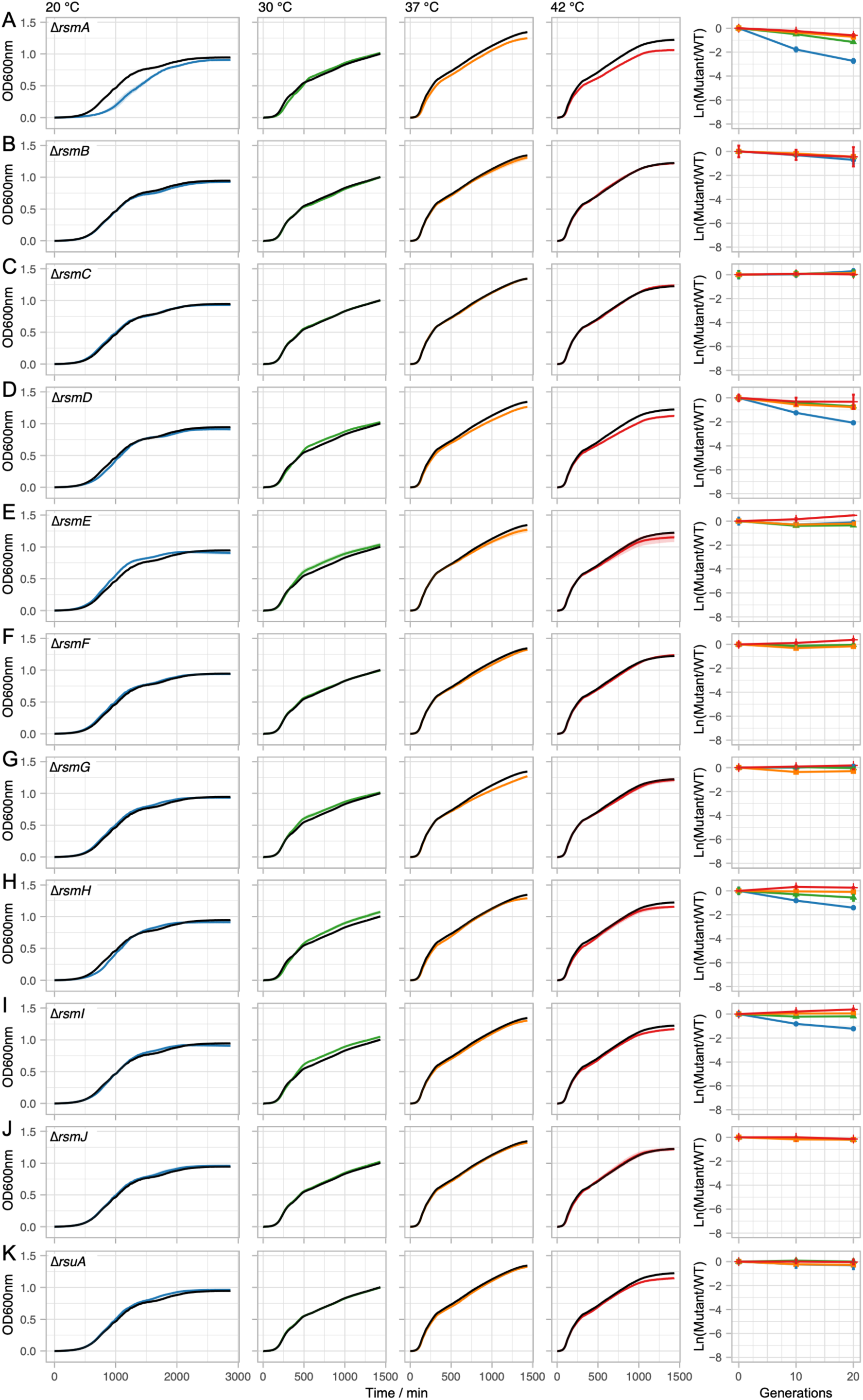
Analysis of growth characteristics of strains with single gene deletions. (**A** - **G**) Growth curves and competition experiments of isolates with unmodified domains and combinations thereof. Strain genotypes are indicated in each panel and growth temperatures are shown at the top of the growth curves. Growth curves of the constructed strains are shown in blue (20 °C), green (30 °C), orange (37 °C), and red (42 °C) and the growth curves of the isogenic wild type at each temperature are shown in black. All growth curves represent the average of three biological replicates. Standard deviation is shown as shaded region. Competitions were performed at 20 °C (blue, circles), 30 °C (green, triangles), 37 °C (orange, squares), and 42 °C (red, line). All competition data represent the average ± standard deviation of three biological replicates. Values were normalized to account for the fitness cost of the fluorescence marker and differences in initial starting ratios of the competitors.

**Table S1.** Overview of modification enzymes and deletions.

| Gene | 16S rRNA modifications<sup>a</sup> | Domain | Deletion (aa) | Deletion (%) |
| --- | --- | --- | --- | --- |
| *rsmA* | m<sup>6</sup><sub>2</sub>A1518, m<sup>6</sup><sub>2</sub>A1519 | 3'm | 7 – 115 | 40 |
| *rsmB* | m<sup>5</sup>C967 | 3'M | 7 – 423 | 97 |
| *rsmC* | m<sup>2</sup>G1207 | 3'M | 7 – 337 | 97 |
| *rsmD* | m<sup>2</sup>G966 | 3'M | 7 – 192 | 94 |
| *rsmE* | m<sup>3</sup>U1498 | 3'm | 7 – 237 | 95 |
| *rsmF* | m<sup>5</sup>C1407, <u>tRNA-Tyr m<sup>5</sup>C49</u> | 3'm | 7 – 473 | 97 |
| *rsmG* | m<sup>7</sup>G527 | 5' | 7 – 201 | 94 |
| *rsmH* | m<sup>4</sup>C1402 | 3'm | 7 – 181 | 56 |
| *rsmI* | Cm1402 | 3'm | 60 – 280 | 77 |
| *rsmJ* | m<sup>2</sup>G1516 | 3'm | 7 – 244 | 95 |
| *rsuA* | Ψ516 | 5' | 7 – 225 | 95 |

<sup>a</sup> Modification of tRNA-Tyr is underlined.

**Table S2.**
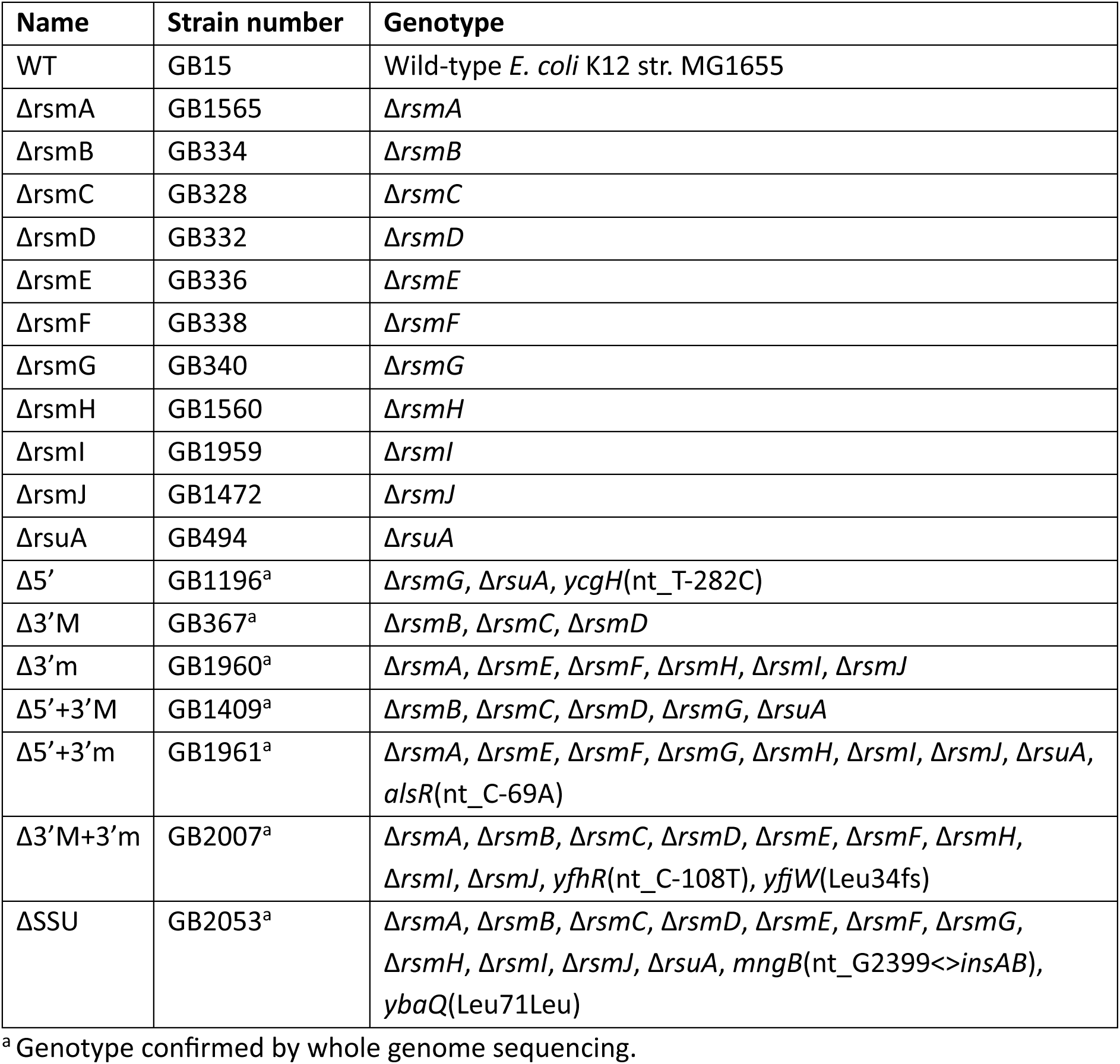
Strain list with relevant genotypes.

**Table S3.** Exponential doubling times of constructed strains.

| Strain | 20 °C |  | 30 °C |  | 37 °C |  | 42 °C |  |
| --- | --- | --- | --- | --- | --- | --- | --- | --- |
|  | Dt ± SD (min) | P-value | Dt ± SD (min) | P-value | Dt ± SD (min) | P-value | Dt ± SD (min) | P-value |
| WT | 112.7 ± 0.5 | - | 34.0 ± 0.3 | - | 21.0 ± 0.2 | - | 18.3 ± 0.3 | - |
| *ΔrsmA* | 165.1 ± 7.5 | 0.007 | 38.5 ± 0.3 | 4.71 × 10<sup>-5</sup> | 21.9 ± 0.4 | 0.045 | 19.2 ± 0.1 | 0.013 |
| *ΔrsmB* | 114.0 ± 0.4 | 0.017 | 34.4 ± 0.3 | 0.182 | 21.1 ± 0.1 | 0.823 | 18.4 ± 0.4 | 0.711 |
| *ΔrsmC* | 108.4 ± 0.6 | 0.001 | 33.6 ± 0.5 | 0.327 | 20.7 ± 0.5 | 0.395 | 18.2 ± 0.3 | 0.768 |
| *ΔrsmD* | 142.6 ± 1.5 | 3.11 × 10<sup>-4</sup> | 35.4 ± 0.4 | 0.011 | 21.2 ± 0.2 | 0.477 | 18.3 ± 0.2 | 1.000 |
| *ΔrsmE* | 116.6 ± 3.3 | 0.175 | 34.0 ± 0.5 | 0.921 | 20.8 ± 0.3 | 0.425 | 18.1 ± 0.2 | 0.422 |
| *ΔrsmF* | 110.6 ± 1.6 | 0.142 | 33.3 ± 0.6 | 0.149 | 20.8 ± 0.1 | 0.182 | 18.3 ± 0.6 | 0.874 |
| *ΔrsmG* | 108.1 ± 1.0 | 0.006 | 33.6 ± 0.3 | 0.222 | 20.7 ± 0.3 | 0.257 | 18.2 ± 0.2 | 0.738 |
| *ΔrsmH* | 144.1 ± 0.4 | 1.06 × 10<sup>-7</sup> | 37.4 ± 0.4 | 0.001 | 21.8 ± 0.1 | 0.012 | 19.0 ± 0.2 | 0.024 |
| *ΔrsmI* | 135.6 ± 0.9 | 4.40 × 10<sup>-5</sup> | 34.2 ± 0.2 | 0.387 | 20.9 ± 0.3 | 0.667 | 18.3 ± 0.1 | 1.000 |
| *ΔrsmJ* | 111.6 ± 0.2 | 0.041 | 34.1 ± 0.2 | 0.769 | 20.8 ± 0.4 | 0.512 | 19.1 ± 1.4 | 0.386 |
| *ΔrsuA* | 109.9 ± 0.6 | 0.004 | 33.3 ± 0.3 | 0.040 | 20.7 ± 0.2 | 0.075 | 18.2 ± 0.1 | 0.566 |
| Δ5' | 117.7 ± 0.4 | 1.42 × 10<sup>-4</sup> | 33.5 ± 0.4 | 0.141 | 20.6 ± 0.2 | 0.090 | 18.5 ± 0.0 | 0.250 |
| Δ3'M | 134.4 ± 2.6 | 0.004 | 35.0 ± 0.1 | 0.020 | 21.0 ± 0.2 | 0.842 | 19.0 ± 0.2 | 0.024 |
| Δ3'm | 244.6 ± 3.9 | 2.46 × 10<sup>-4</sup> | 52.4 ± 0.3 | 2.29 × 10<sup>-7</sup> | 24.7 ± 0.3 | 2.22 × 10<sup>-4</sup> | 20.4 ± 0.2 | 0.001 |
| Δ5'+3'M | 131.9 ± 4.3 | 0.016 | 34.9 ± 0.5 | 0.079 | 21.6 ± 0.2 | 0.024 | 19.6 ± 0.3 | 0.003 |
| Δ5'+3'm | 219.0 ± 1.9 | 4.51 × 10<sup>-5</sup> | 54.6 ± 0.2 | 3.09 × 10<sup>-7</sup> | 28.8 ± 0.4 | 3.18 × 10<sup>-5</sup> | 23.1 ± 0.1 | 1.99 × 10<sup>-4</sup> |
| Δ3'M+3'm | 306.0 ± 3.2 | 6.94 × 10<sup>-5</sup> | 53.3 ± 0.2 | 2.49 × 10<sup>-6</sup> | 28.3 ± 0.2 | 1.76 × 10<sup>-6</sup> | 25.3 ± 0.1 | 4.47 × 10<sup>-5</sup> |
| ΔSSU | 313.7 ± 6.0 | 2.72 × 10<sup>-4</sup> | 56.8 ± 0.7 | 3.91 × 10<sup>-5</sup> | 31.6 ± 0.3 | 8.96 × 10<sup>-7</sup> | 28.5 ± 0.2 | 3.90 × 10<sup>-6</sup> |

**Table S4.** Relative fitness^a^ during exponential growth of constructed strains.

| Strain | 20 °C |  | 30 °C |  | 37 °C |  | 42 °C |  |
| --- | --- | --- | --- | --- | --- | --- | --- | --- |
|  | Fitness ± SD | P-value | Fitness ± SD | P-value | Fitness ± SD | P-value | Fitness ± SD | P-value |
| WT | 1.00 ± 0.00 | - | 1.00 ± 0.01 | - | 1.00 ± 0.01 | - | 1.00 ± 0.01 | - |
| *ΔrsmA* | 0.68 ± 0.03 | 0.007 | 0.88 ± 0.01 | 4.71 × 10<sup>-5</sup> | 0.96 ± 0.02 | 0.045 | 0.95 ± 0.01 | 0.013 |
| *ΔrsmB* | 0.99 ± 0.00 | 0.017 | 0.99 ± 0.01 | 0.182 | 1.00 ± 0.01 | 0.823 | 1.00 ± 0.02 | 0.711 |
| *ΔrsmC* | 1.04 ± 0.01 | 0.001 | 1.01 ± 0.02 | 0.327 | 1.01 ± 0.03 | 0.395 | 1.01 ± 0.01 | 0.768 |
| *ΔrsmD* | 0.79 ± 0.01 | 3.11 × 10<sup>-4</sup> | 0.96 ± 0.01 | 0.011 | 0.99 ± 0.01 | 0.477 | 1.00 ± 0.01 | 1.000 |
| *ΔrsmE* | 0.97 ± 0.03 | 0.175 | 1.00 ± 0.01 | 0.921 | 1.01 ± 0.02 | 0.425 | 1.01 ± 0.01 | 0.422 |
| *ΔrsmF* | 1.02 ± 0.01 | 0.142 | 1.02 ± 0.02 | 0.149 | 1.01 ± 0.00 | 0.182 | 1.00 ± 0.03 | 0.874 |
| *ΔrsmG* | 1.04 ± 0.01 | 0.006 | 1.01 ± 0.01 | 0.222 | 1.01 ± 0.02 | 0.257 | 1.01 ± 0.01 | 0.738 |
| *ΔrsmH* | 0.78 ± 0.00 | 1.06 × 10<sup>-7</sup> | 0.91 ± 0.01 | 0.001 | 0.96 ± 0.00 | 0.012 | 0.96 ± 0.01 | 0.024 |
| *ΔrsmI* | 0.83 ± 0.01 | 4.40 × 10<sup>-5</sup> | 0.99 ± 0.01 | 0.387 | 1.00 ± 0.01 | 0.667 | 1.00 ± 0.00 | 1.000 |
| *ΔrsmJ* | 1.01 ± 0.00 | 0.041 | 1.00 ± 0.01 | 0.769 | 1.01 ± 0.02 | 0.512 | 0.96 ± 0.07 | 0.386 |
| *ΔrsuA* | 1.03 ± 0.01 | 0.004 | 1.02 ± 0.01 | 0.040 | 1.02 ± 0.01 | 0.075 | 1.01 ± 0.00 | 0.566 |
| Δ5' | 0.96 ± 0.00 | 1.42 × 10<sup>-4</sup> | 1.02 ± 0.01 | 0.141 | 1.02 ± 0.01 | 0.090 | 0.99 ± 0.00 | 0.250 |
| Δ3'M | 0.84 ± 0.02 | 0.004 | 0.97 ± 0.00 | 0.020 | 1.00 ± 0.01 | 0.842 | 0.96 ± 0.01 | 0.024 |
| Δ3'm | 0.46 ± 0.01 | 2.46 × 10<sup>-4</sup> | 0.65 ± 0.00 | 2.29 × 10<sup>-7</sup> | 0.85 ± 0.01 | 2.22 × 10<sup>-4</sup> | 0.90 ± 0.01 | 0.001 |
| Δ5'+3'M | 0.86 ± 0.03 | 0.016 | 0.97 ± 0.01 | 0.079 | 0.97 ± 0.01 | 0.024 | 0.93 ± 0.01 | 0.003 |
| Δ5'+3'm | 0.51 ± 0.00 | 4.51 × 10<sup>-5</sup> | 0.62 ± 0.00 | 3.09 × 10<sup>-7</sup> | 0.73 ± 0.01 | 3.18 × 10<sup>-5</sup> | 0.79 ± 0.00 | 1.99 × 10<sup>-4</sup> |
| Δ3'M+3'm | 0.37 ± 0.00 | 6.94 × 10<sup>-5</sup> | 0.64 ± 0.00 | 2.49 × 10<sup>-6</sup> | 0.74 ± 0.01 | 1.76 × 10<sup>-6</sup> | 0.72 ± 0.00 | 4.47 × 10<sup>-5</sup> |
| ΔSSU | 0.36 ± 0.01 | 2.72 × 10<sup>-4</sup> | 0.60 ± 0.01 | 3.91 × 10<sup>-5</sup> | 0.66 ± 0.01 | 8.96 × 10<sup>-7</sup> | 0.64 ± 0.00 | 3.90 × 10<sup>-6</sup> |

<sup>a</sup> Growth fitness relative to the isogenic wild type at each respective temperature. See Supplementary Table S3 for doubling times.

**Table S5.** Competitive fitness^a^ during exponential growth of constructed strains.

| Strain | 20 °C |  | 30 °C |  | 37 °C |  | 42 °C |  |
| --- | --- | --- | --- | --- | --- | --- | --- | --- |
|  | Fitness ± SD | P-value | Fitness ± SD | P-value | Fitness ± SD | P-value | Fitness ± SD | P-value |
| WT | 1.00 ± 0.00 | - | 1.00 ± 0.00 | - | 1.00 ± 0.01 | - | 1.00 ± 0.01 | - |
| *ΔrsmA* | 0.86 ± 0.01 | 3.46 x 10<sup>-4</sup> | 0.94 ± 0.00 | 2.05 x 10<sup>-4</sup> | 0.96 ± 0.01 | 0.010 | 0.97 ± 0.00 | 0.005 |
| *ΔrsmB* | 0.96 ± 0.01 | 0.031 | 0.98 ± 0.01 | 0.018 | 0.98 ± 0.00 | 0.060 | 0.98 ± 0.02 | 0.121 |
| *ΔrsmC* | 1.01 ± 0.01 | 0.059 | 1.01 ± 0.00 | 0.084 | 1.01 ± 0.00 | 0.403 | 1.00 ± 0.00 | 0.829 |
| *ΔrsmD* | 0.90 ± 0.00 | 1.04 x 10<sup>-5</sup> | 0.96 ± 0.00 | 2.19 x 10<sup>-4</sup> | 0.96 ± 0.00 | 0.017 | 0.98 ± 0.02 | 0.242 |
| *ΔrsmE* | 0.99 ± 0.01 | 0.497 | 0.98 ± 0.00 | 0.004 | 0.99 ± 0.00 | 0.215 | 1.03 ± 0.00 | 0.013 |
| *ΔrsmF* | 0.99 ± 0.00 | 0.019 | 1.00 ± 0.00 | 0.459 | 0.99 ± 0.00 | 0.307 | 1.02 ± 0.00 | 0.020 |
| *ΔrsmG* | 1.01 ± 0.00 | 0.052 | 1.00 ± 0.00 | 0.385 | 0.99 ± 0.00 | 0.121 | 1.01 ± 0.01 | 0.341 |
| *ΔrsmH* | 0.93 ± 0.01 | 0.002 | 0.97 ± 0.01 | 0.004 | 1.00 ± 0.00 | 0.537 | 1.01 ± 0.00 | 0.065 |
| *ΔrsmI* | 0.94 ± 0.01 | 0.001 | 0.99 ± 0.00 | 0.032 | 1.00 ± 0.00 | 0.672 | 1.02 ± 0.00 | 0.020 |
| *ΔrsmJ* | 0.99 ± 0.01 | 0.058 | 0.99 ± 0.00 | 0.026 | 0.99 ± 0.00 | 0.237 | 0.99 ± 0.00 | 0.202 |
| *ΔrsuA* | 0.98 ± 0.01 | 0.085 | 1.00 ± 0.00 | 0.930 | 0.99 ± 0.00 | 0.140 | 1.00 ± 0.00 | 0.723 |
| *Δ5'* | 0.99 ± 0.00 | 0.004 | 1.00 ± 0.00 | 0.487 | 0.99 ± 0.00 | 0.337 | 1.01 ± 0.00 | 0.101 |
| *Δ3'M* | 0.89 ± 0.00 | 4.91 x 10<sup>-7</sup> | 0.95 ± 0.00 | 4.53 x 10<sup>-4</sup> | 0.95 ± 0.00 | 0.010 | 0.96 ± 0.00 | 0.003 |
| *Δ3'm* | 0.70 ± 0.01 | 0.001 | 0.84 ± 0.00 | 1.30 x 10<sup>-6</sup> | 0.89 ± 0.01 | 4.50 x 10<sup>-4</sup> | 0.93 ± 0.00 | 4.78 x 10<sup>-4</sup> |
| *Δ5'+3'M* | 0.91 ± 0.00 | 1.53 x 10<sup>-6</sup> | 0.96 ± 0.00 | 0.001 | 0.95 ± 0.00 | 0.015 | 0.97 ± 0.01 | 0.007 |
| *Δ5'+3'm* | 0.64 ± 0.01 | 5.67 x 10<sup>-6</sup> | 0.80 ± 0.00 | 6.81 x 10<sup>-7</sup> | 0.80 ± 0.01 | 6.96 x 10<sup>-5</sup> | 0.86 ± 0.01 | 0.001 |
| *Δ3'M+3'm* | 0.52 ± 0.01 | 9.57 x 10<sup>-5</sup> | 0.75 ± 0.01 | 6.11 x 10<sup>-6</sup> | 0.71 ± 0.00 | 8.34 x 10<sup>-5</sup> | 0.72 ± 0.01 | 1.29 x 10<sup>-6</sup> |
| *ΔSSU* | 0.50 ± 0.01 | 1.27 x 10<sup>-5</sup> | 0.71 ± 0.01 | 2.13 x 10<sup>-6</sup> | 0.67 ± 0.02 | 1.36 x 10<sup>-4</sup> | 0.69 ± 0.01 | 9.01 x 10<sup>-7</sup> |

<sup>a</sup> Fitness calculated based on competition experiments relative to the isogenic wild type.

**Table S6.**
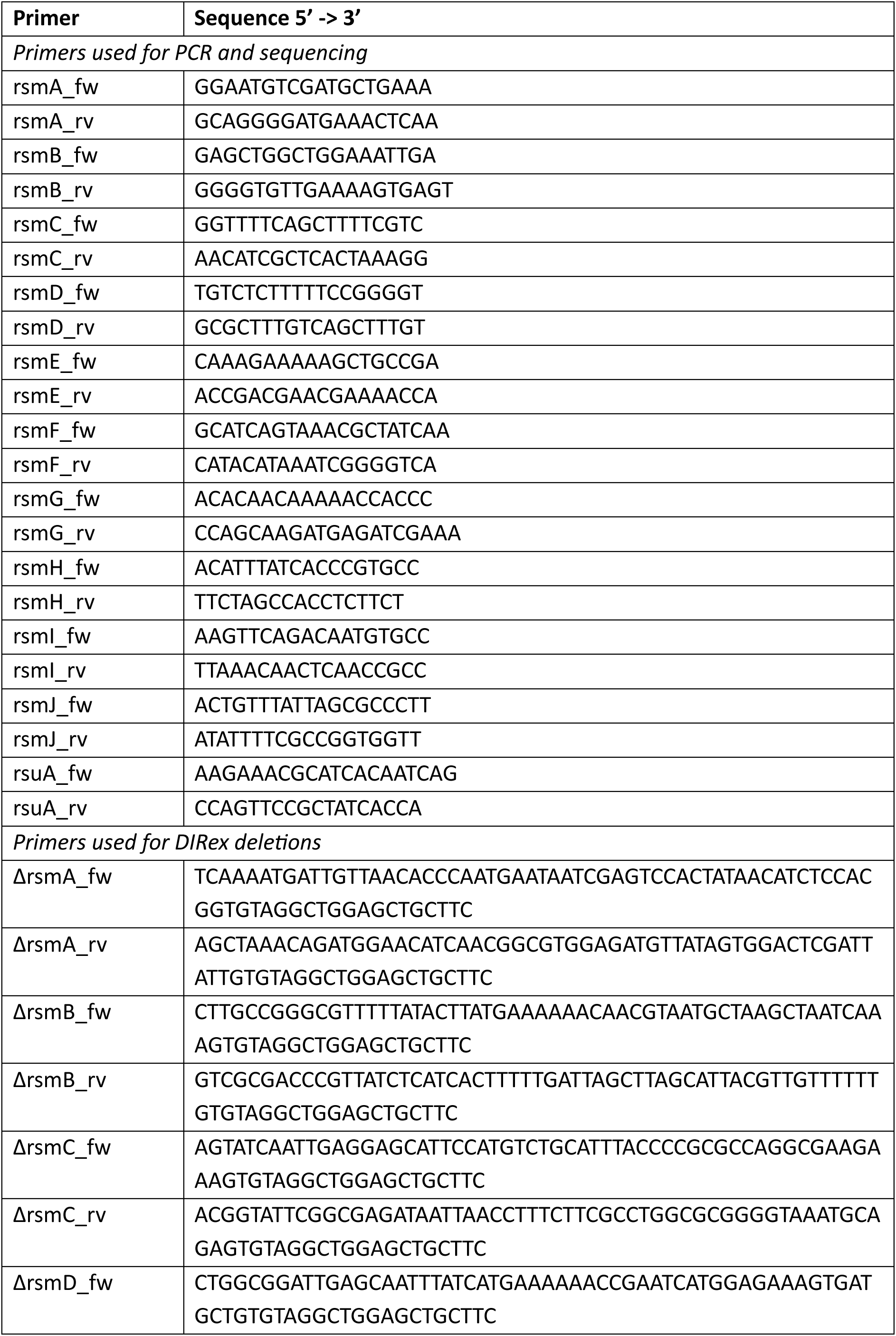
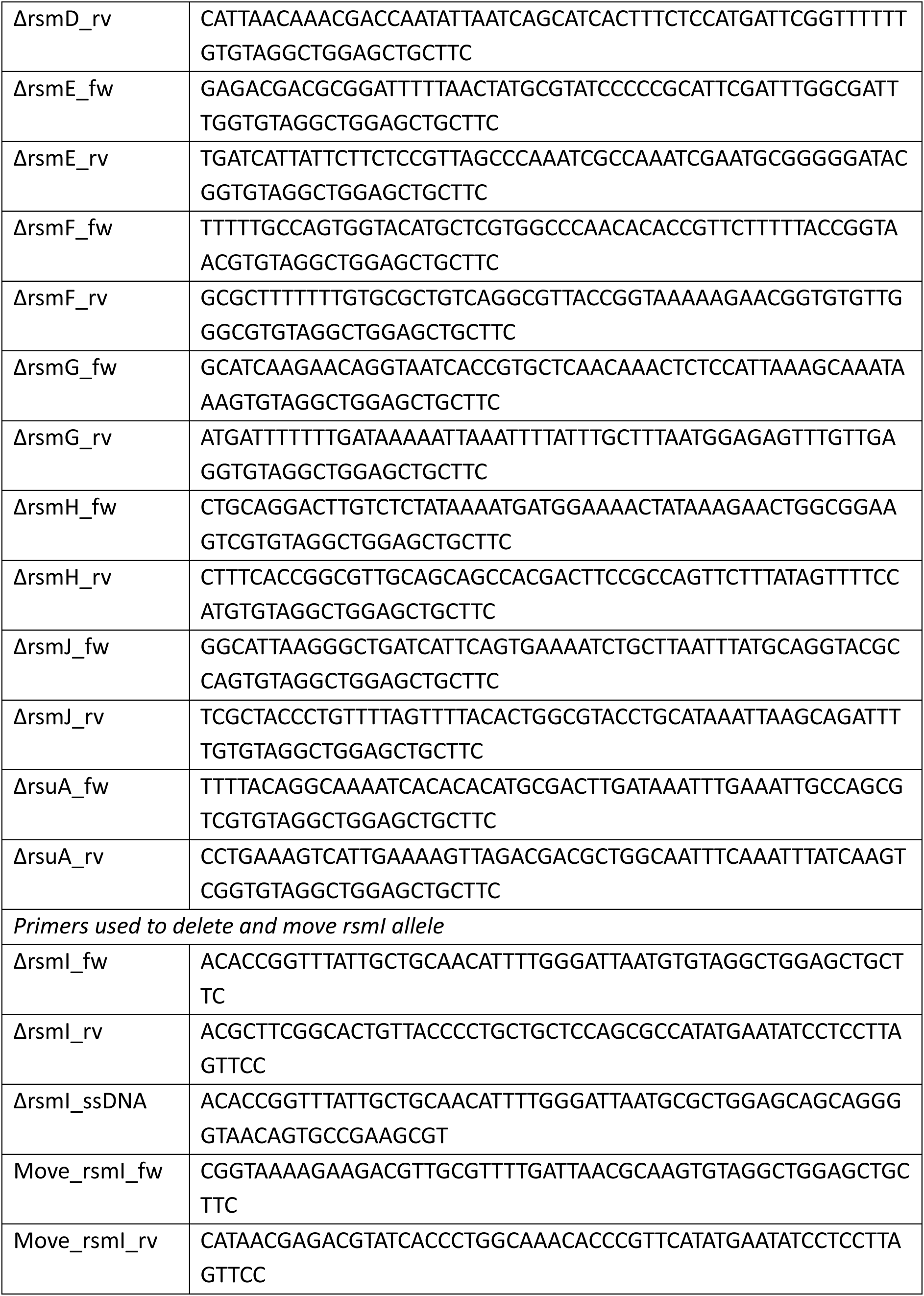
List of oligonucleotides.

**Table S7.** List of antibiotic disc tests.

| **Antibiotic** | **Quantity (µg)** |
| --- | --- |
| Kasugamycin | 300 |
| Gentamicin | 30 |
| Streptomycin | 300 |
| Amikacin | 30 |
| Spectinomycin | 25 |
| Tetracyclin | 30 |
| Tigecyclin | 15 |
| Linezolid | 30 |
| Chloramphenicol | 30 |
| Erythromycin | 30 |
| Clindamycin | 10 |
| Tiamulin | 30 |
| Rifampicin | 30 |

